# A novel sorting method reveals superior metabolic activity of mononucleated over binucleated tetraploid hepatocytes

**DOI:** 10.1101/2025.05.07.652591

**Authors:** Yusuke Watanabe, Jonathan H. Sussman, Tomoya Anami, Moena Hattori, Saaya Yamane, Masaki Nishikawa, Yasuyuki Sakai, Takeshi Katsuda

## Abstract

Hepatocytes display notable ploidy diversity, varying both in the number of genomic copies and in the number of nuclei. Most mouse hepatocytes are tetraploid (4n), which can exist as either mononucleated (1×4n) or binucleated (2×2n) cells. Despite this distinction, these two cell types have traditionally been grouped and studied as a single population. One likely reason for this is that conventional ploidy-sorting methods classify hepatocytes based solely on total DNA content, without distinguishing between mononucleated and binucleated states. In this study, we developed a novel flow cytometry strategy to distinguish and isolate 1×4n and 2×2n hepatocytes. Our approach leverages Hoechst-Area to assess total ploidy and Hoechst-Height to differentiate mononucleated and binucleated hepatocytes. Transcriptome analysis comparing these two populations revealed that 1×4n hepatocytes exhibit a broader and more metabolically active gene expression profile. Importantly, this enhanced metabolic activity was independent of liver zonation, a well-known driver of metabolic heterogeneity in hepatocytes. Our findings uncover a previously underappreciated layer of functional diversity in the liver and provide a new framework for studying the physiological and pathological roles of nuclear configuration in hepatocytes.

## MAIN

Unlike most other mammalian cell types, hepatocytes exhibit variable numbers of genomic copies (ploidy). The most common state in mouse hepatocytes is a binucleated tetraploid cell with each nucleus containing two genomic copies (2×2n), while other cells are mononucleated with four genomic copies (1×4n). A smaller percentage of hepatocytes contain two tetraploid nuclei (2x4n) or one nucleus with 8 genome copies (1×8n). While tetraploid hepatocytes have traditionally been studied as a single entity, several recent studies reported that binucleated hepatocytes decrease in pathological conditions such as after a partial hepatectomy[1] and carbon tetrachloride (CCl_4_)-induced injury[2], suggesting that there are functional differences between mononucleated and binucleated hepatocytes.

A major limitation in studying the differences between 1×4n and 2×2n hepatocytes has been the lack of a method to isolate them. A flow cytometry study in plants reported that DNA-stained pollen populations could be distinguished based on their fluorescence and intensity parameters when using propidium iodide (PI), allowing for selective sorting of mononucleated and binucleated populations[3]. We hypothesized that a similar approach could be applied to hepatocytes, using Hoechst 33342, a DNA dye compatible with live cells. First, we confirmed that the conventional gating strategy enabled the sorting of freshly isolated mouse hepatocytes with different ploidy states (**Figure 1A, Supplementary Figure S1**), yielding mononucleated or binucleated cells whose nuclear sizes increased in accordance with their expected ploidy states (**Figure 1B**). We then analyzed hepatocytes using Hoechst-Area, representing the total integrated fluorescence signal, and Hoechst-Height, representing the maximum intensity of the signal pulse. As expected, 4n hepatocytes separated into two distinct populations, one with a higher Hoechst-Height signal (termed 4n-top) and one with a lower signal (termed 4n-bottom) (**Figure 1C**). To examine whether these populations correspond to 1×4n and 2×2n hepatocytes, we sorted the whole 4n (4n-whole), 4n-top and 4n-bottom hepatocytes. The proportions of mononucleated and binucleated cells were determined by microscopy after plating, confirming that 4n-top and 4n-bottom populations were enriched for mononucleated and binucleated cells, respectively (**Figure 1D-E**). Specifically, the 4n-whole population consisted of 48.0% of mononucleated and 52.0 ± 3.7% of binucleated cells (mean ± SEM). In contrast, the mononucleated fraction in the 4n-top population increased to 59.6 ± 1.6% while the binucleated fraction in the 4n-bottom population reached 87.5 ± 2.1% of 2×2n cells (n = 9) (**Figure 1E**). To further test our hypothesis that the 4n-top population is enriched for 1×4n hepatocytes, we performed three rounds of sequential sorting. This gradually increased the proportion of 1×4n cells to 75.5 ± 1.2% (n = 3) (**Figure 1F-G**). In conclusion, our new gating strategy enables the selective sorting of the 1×4n and 2×2n hepatocytes.

**Figure 1.**
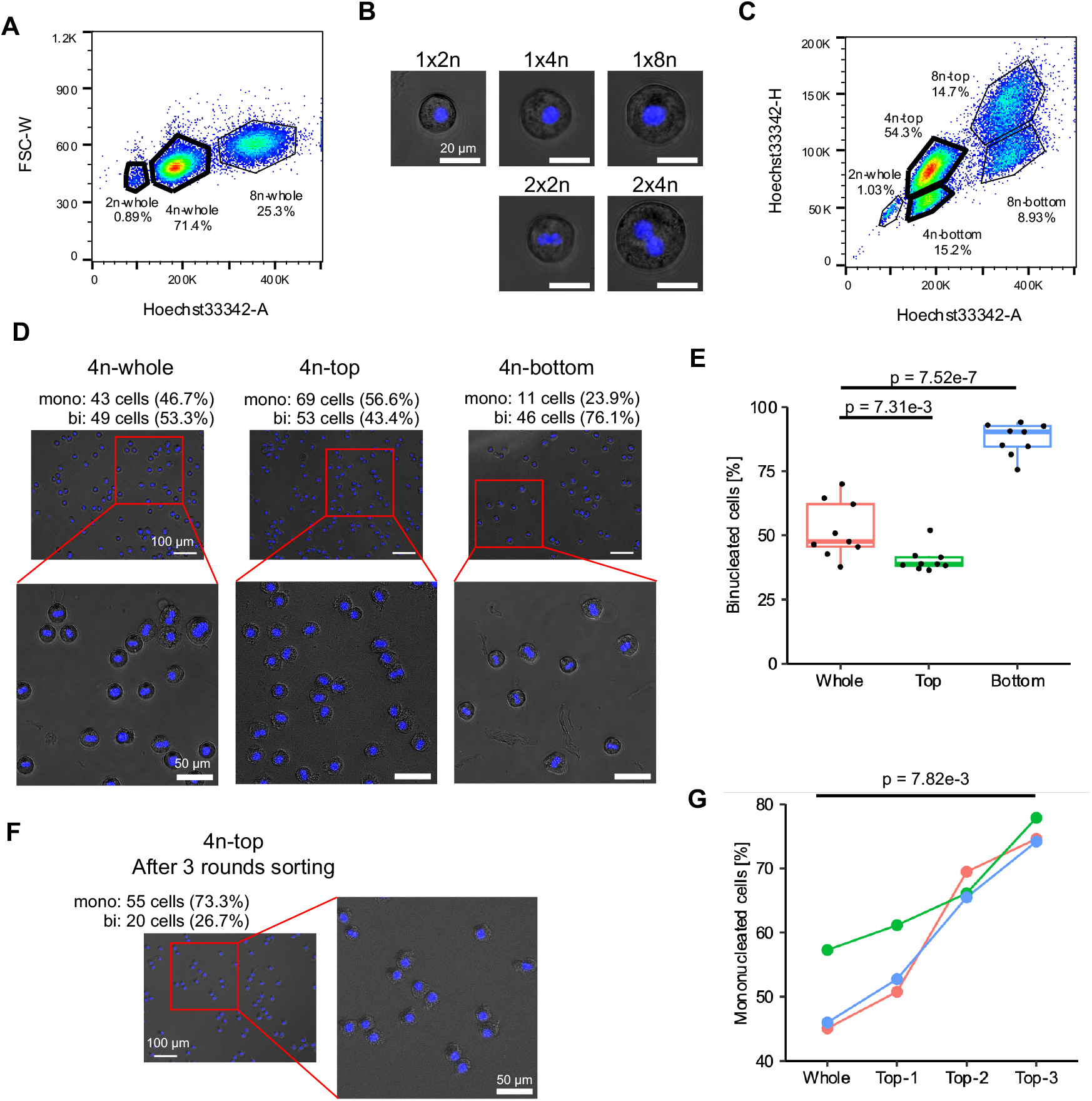
Development of a novel strategy for isolating 1×4n and 2×2n hepatocytes. (A) The conventional gating strategy for sorting whole 2n, 4n and 8n hepatocytes. (B) Representative images of sorted hepatocytes with distinct ploidy states. (C) The newly developed gating strategy for presumable 1×4n and 2×2n hepatocyte populations. (D) Representative images of plated 4n hepatocytes sorted with the conventional (**A**) or new (**C**) gating strategy. (E) Proportions of binucleated cells in each of the sorted 4n populations (n = 9). Significance was assessed using a one-sided Welch’s *t-*test. (F) A representative image of plated cells after three rounds of sorting of 4n-top hepatocytes. (G) Proportions of mononucleated cells after three rounds of sorting of 4n-top hepatocytes (n = 3). Significance was assessed using a one-sided Welch’s t-test.

Using this sorting method, we next investigated differences of gene expression profiles among hepatocytes with different ploidy. We sorted 2n-whole, 4n-whole, 4n-top and 4n-bottom populations and performed RNA sequencing (RNA-seq). For 4n-top hepatocytes, we used cells after a single round of sorting, rather than repeated sorting, to ensure enough cells for RNA-seq. We first performed a global principal component analysis (PCA) across all the samples, which revealed a clear separation between the 2n-whole population and the three 4n populations along the first principal component (PC1) (**Figure 2A**). A binary comparison of the 4n-whole versus the 2n-whole population identified 1363 differentially expressed genes (DEGs, p-adj<0.05, |log2FC|>0.5) (**Figure 2B**) and 934 differentially enriched pathways (FDR<0.05) from the Gene Ontology Biological Process (GOBP) category using a Gene Set Enrichment Analysis (GSEA) (top enriched pathways shown in **Figure 2C**). These results indicated that our RNA-seq dataset reliably captures the difference between 2n and 4n cells, while representing the overall similarity among the 4n populations.

**Figure 2.**
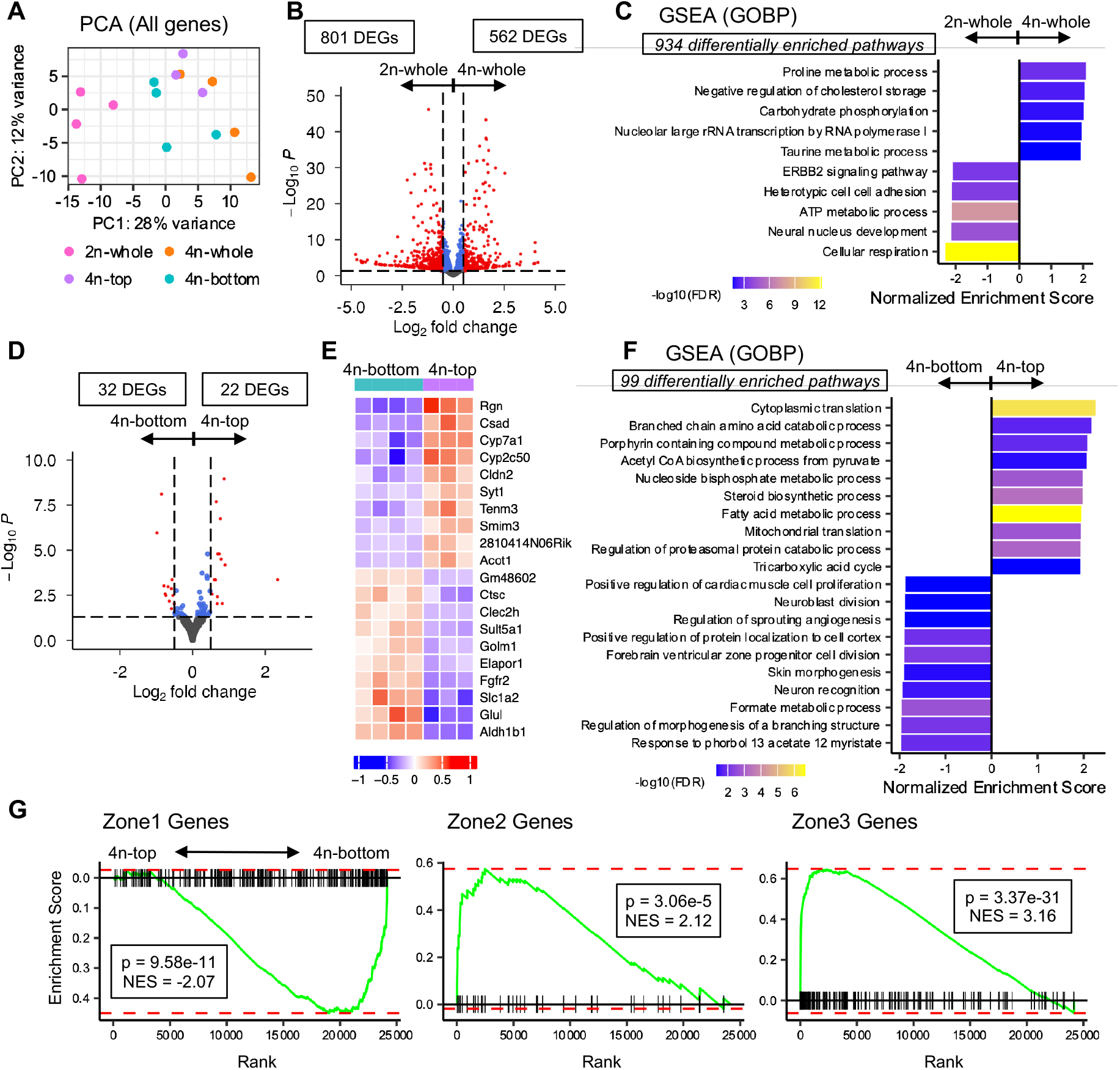
Transcriptomic analysis of hepatocytes with different ploidy states. (A) Principal component analysis (PCA) across all samples. (B) A volcano plot comparing gene expression between 4n-whole and 2n-whole hepatocytes. The log fold change shrunken data was plotted. Significant genes defined as p-adj < 0.05 and |Log2FC| > 0.5. (c) Representative pathways differentially enriched between 4n-whole and 2n-whole hepatocytes. The Gene Ontology Biological Process (GOBP) database was used. FDR, false discovery rate. (D) A volcano plot comparing 4n-top and 4n-bottom hepatocytes. The log fold change shrunken data was plotted. Significant genes defined as p-adj < 0.05 and |Log2FC| > 0.5. (E) A heatmap comparing 4n-top and 4n-bottom hepatocytes. Among the 54 DEGs, the top 10 genes enriched in 4n-top and 4n-bottom are shown, respectively. Genes are colored by z-score normalized expression. (F) Representative pathways differentially enriched between 4n-top and 4n-bottom hepatocytes. (G) GSEA using manually curated gene lists characteristic to each zone. The analysis was conducted by comparing 4n-top (left) and 4n-bottom (right) hepatocytes. FDR, false discovery rate. NES, normalized enrichment score.

We next investigated differences between 1×4n and 2×2n hepatocytes. Notably, the 4n-bottom population was positioned closer to the 2n-whole population along PC1 than the other 4n populations were. This observation is consistent with a previous report suggesting that the gene expression profile of 2×2n nuclei is more similar to that of 1×2n nuclei than 1×4n nuclei[4]. Despite the overall similarity, a direct comparison between 4n-top and 4n-bottom populations identified 54 DEGs (**Figure 2D**, representative genes shown in **Figure 2E**). Moreover, GSEA revealed differential enrichment across multiple pathway categories, including 99 in the GOBP (top enriched pathways shown in **Figure 2F**), 14 in KEGG, 122 in Reactome and 10 in Hallmark (**Supplementary Figure S2A**). Notably, many pathways enriched in 4n-top were involved in metabolism, including *branched chain amino acid catabolism* and *porphyrin-containing compound metabolism*. Given that the liver is spatially portioned into three metabolic zones (known as zonation), we considered whether these differences were driven by zonation rather than ploidy. To assess the potential influence of zonation, we curated zone-specific gene lists based on published RNA-seq datasets[5,6] GSEA using these lists revealed that Zone 1 genes were enriched in 4n-bottom, while Zone 2 and 3 genes were enriched in 4n-top (**Figure 2G**). This result suggests that 2×2n hepatocytes are more prevalent in Zone 1, whereas 1×4n hepatocytes are enriched in Zone 2 and 3, consistent with previous reports[4,7].

To further explore the relationship between ploidy and zonation, we examined enrichment of zonation-associated metabolic pathways[8,9]. Unexpectedly, Zone 1-related *fatty acid β-oxidation* and Zone 3-related *xenobiotic metabolism* were both enriched in 4n-top hepatocytes (**Supplementary Figure S2B**). Moreover, 4n-top showed enrichment of many other metabolism-related pathways that were not previously linked to zonation, including *nucleoside bisphosphate metabolism* and *tetrapyrrole metabolism* (**Supplementary Figure S2C**). In contrast, we found only two metabolic pathways enriched in the 4n-bottom population: *formate metabolism* and *glutamine family amino acid metabolism* (**Supplementary Figure S2D**). Taken together, these analyses suggest that zonation alone cannot fully explain the observed differences between 1×4n and 2×2n hepatocytes. Therefore, it is likely that these two populations have inherently distinct metabolic programs, with 1×4n hepatocytes exhibiting broader and more active metabolic profiles.

In summary, our novel sorting strategy enabled the selective isolation of 1×4n and 2×2n hepatocytes, allowing us to uncover transcriptomic differences between these two populations. Although the greatest differences existed between 4n and 2n cells, 1×4n and 2×2n hepatocytes exhibited clear transcriptional differences, including more active metabolic profiles in 1×4n hepatocytes compared to 2×2n hepatocytes. These findings motivate future work on the roles of different nucleation states on cellular function in the liver. Recent developments in transcriptional profiling, such as spatial transcriptomics will facilitate a more granular dissection of the phenotypes contained within each ploidy state.

## METHODS

### Mice

C57BL/6JJcl mice were purchased from CLEA Japan. Mice were fed with a standard diet CE-2 (10 kGy) (CLEA Japan) and kept in a 12-hour light–dark cycle. 7- to 15-week-old male mice were used for experiments.

### Hepatocyte isolation

Hepatocyte isolation from the mouse liver was performed as previously described[10]. Briefly, livers were perfused with HBSS (Wako), Liver Perfusion Medium (Gibco), and HBSS with 5 mM CaCl2 (Wako) and 500 μg/ml collagenase (Wako) in order. Following perfusion, livers were mechanically dispersed with tweezers, resuspended in wash medium (DMEM (Wako) with 5% FBS (BioWest)) supplemented with 40 μg/ml DNase (Millipore), and filtrated with a 70 μm cell strainer. The cells were centrifuged at 4 °C at 48 × g for 5 min. Then, the cells were resuspended in complete percoll solution (10.8 ml percoll (Cytiva), 12.5 ml wash medium, and 1.2 ml 10× HBSS per liver) and centrifuged at 4 °C at 48 × g for 10 min. After a single wash with wash medium, hepatocytes and were use d for the downstream experiments.

### Fluorescent staining

Hepatocytes were suspended with DMEM supplemented with 10% FBS and 1 x Penicillin-Streptomycin-Amphotericin B Suspension (P.S.A.) (Wako) at a concentration of 2e6 /ml. After 15 μg/ml Hoechst 33342 (Wako), 5 μM reserpine (Wako) and 40 μg/ml DNase were added, hepatocytes were incubated at 37 °C for 30 min, with inverted every 10 minutes. Following the centrifuge at 4 °C at 48 × g for 5 min, hepatocytes were resuspended with flow buffer (HBSS supplemented with 1 M HEPES (Sigma), 1 M MgCl_2_(Wako), 200 mg/ml glucose (Sigma), 1 x P.S.A., 1 x MEM Non-essential Amino Acids Solution (Wako), 1 X GlutaMAX Supplement (Wako), 1 mM sodium pyruvate (Wako), 40 μg/ml DNase, 15 μg/ml Hoechst 33342 and 5 % FBS and pH was adjusted to 7.4 with NaOH (Wako)) at a concentration of 1e7 /ml. Then, hepatocytes were filtrated with a 40 μm cell strainer, and 100 nm/ml TO-PRO-3 Iodide (Invitrogen) was added for staining dead cells.

### Flow cytometry

Flow cytometry was performed using SH800S (Sony) with a 130 μm nozzle. Events other than hepatocytes and doublets were removed by gating on BSC-A vs. FSC-A and FSC-H vs. FSC-A, respectively. Dead cells were removed according to TO-PRO-3 fluorescence by gating on FSC-H vs. APC-A. Then, hepatocytes were plotted by FSC-W vs. Hoechst 33342-A or Hoechst 33342-H vs. Hoechst 33342-A. Sorted cells were collected into 1:1 mixture of DMEM and FBS. After sorting, cells were immediately centrifuged at 4 °C at 48 × g for 5 min for 4n and 8n cells, or 800 × g for 10 min for 2n cells, respectively. Cells were then resuspended with DMEM supplemented with 10% FBS and 1 x P.S.A. and kept on ice before the downstream experiments.

### Counting of mononucleated and binucleated hepatocytes

A portion of sorted hepatocytes were plated onto a 96-well plate with DMEM supplemented with 10% FBS and 1 x P.S.A. Cells were incubated at 37 °C to for a few hours, stained with 1 μg/ml Hoechst 33342 again and observed by fluorescence microscope BZ-X810 (Keyence). The numbers of mononucleated and binucleated cells were counted, respectively, and the proportions of these cells in sorted samples were calculated.

### RNA extraction

80 ∼ 300 K of hepatocytes were sorted by flowcytometry. RNA was extracted with FastGene RNA basic kit including DNase I treatment. The concentration, A260/A280 and A260/A230 of extracted RNA were measured with NanoDrop One (Thermo). RNA with low purity was cleaned up with Monarch Spin RNA Cleanup Kit (10 μg) (New England Biolabs).

### RNA-seq

Library preparation and sequencing were performed by Azenta (Tokyo, Japan) using NovaSeq (Illumina).

### Bioinformatics for RNA-seq

Reads were aligned to the mouse genome (GRCm39) using STAR aligner with default parameters[11]. The gene-count matrix was produced by featureCounts[12]. Batch effects due to individual differences among mice were adjusted using ComBat. Then, to compare gene expression between samples, expression levels were normalized on the “median of ratios” method using DESeq2[12]. Binary comparisons were performed using the results() function in DEseq2. Differentially expressed genes were defined following the criteria: adjusted p value < 0.05 and |log_2_(fold-change)| > 0.5. Differentially expressed pathways were identified by gene sets enrichment analysis with the cut-off values of false discovery rate < 0.05.

### Preparation of lists of zonated genes

For Zone1- and 3-related genes, the counts per million matrix from PRJNA556572[5] was used. Eight layers along hepatic zones were merged in pairs to create the expression values for four layers, and Kruskal-Walis test was performed for every gene. 188 of Zone1- and 259 of Zone3-related genes were identified by two criteria: FDR < 0.05 and the median values of the layers exhibit a monotonic increase or decrease. For Zone2-related genes, we used 46 of previously identified genes by Halpern *et al*[6].

## ACKNOWLEDGEMENTS

This work was supported by grants from the Uehara Memorial Foundation and the Nakatomi Foundation.

## SUPPLEMENTARY FIGURES

**Supplementary Figure S1. Flow cytometry gating strategy**.

**Supplementary Figure S2. Additional GSEA results**.

(A) Top enriched pathways by GSEA using KEGG, Hallmark, and Reactome categories.

(B-D) GSEA plots comparing 4n-top (left) and 4n-bottom (right). Zonated pathways (B). pathways enriched in 4n-top (C), and pathways enriched in 4n-bottom (D).

