## Supplemental Figures for "A novel sorting method reveals superior metabolic activity of mononucleated over binucleated tetraploid hepatocytes"

### Slide 1
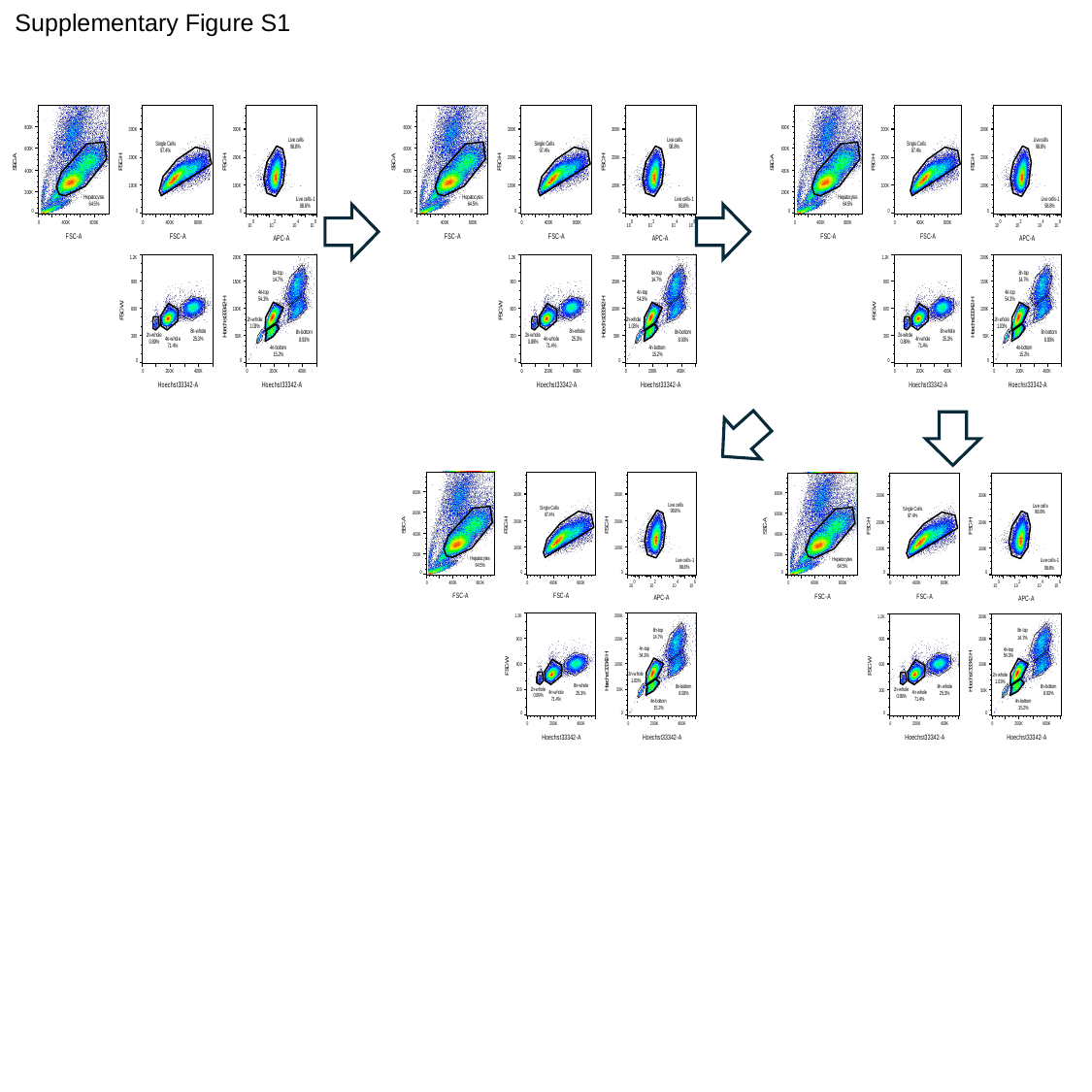

Supplementary Figure S1

### Slide 2
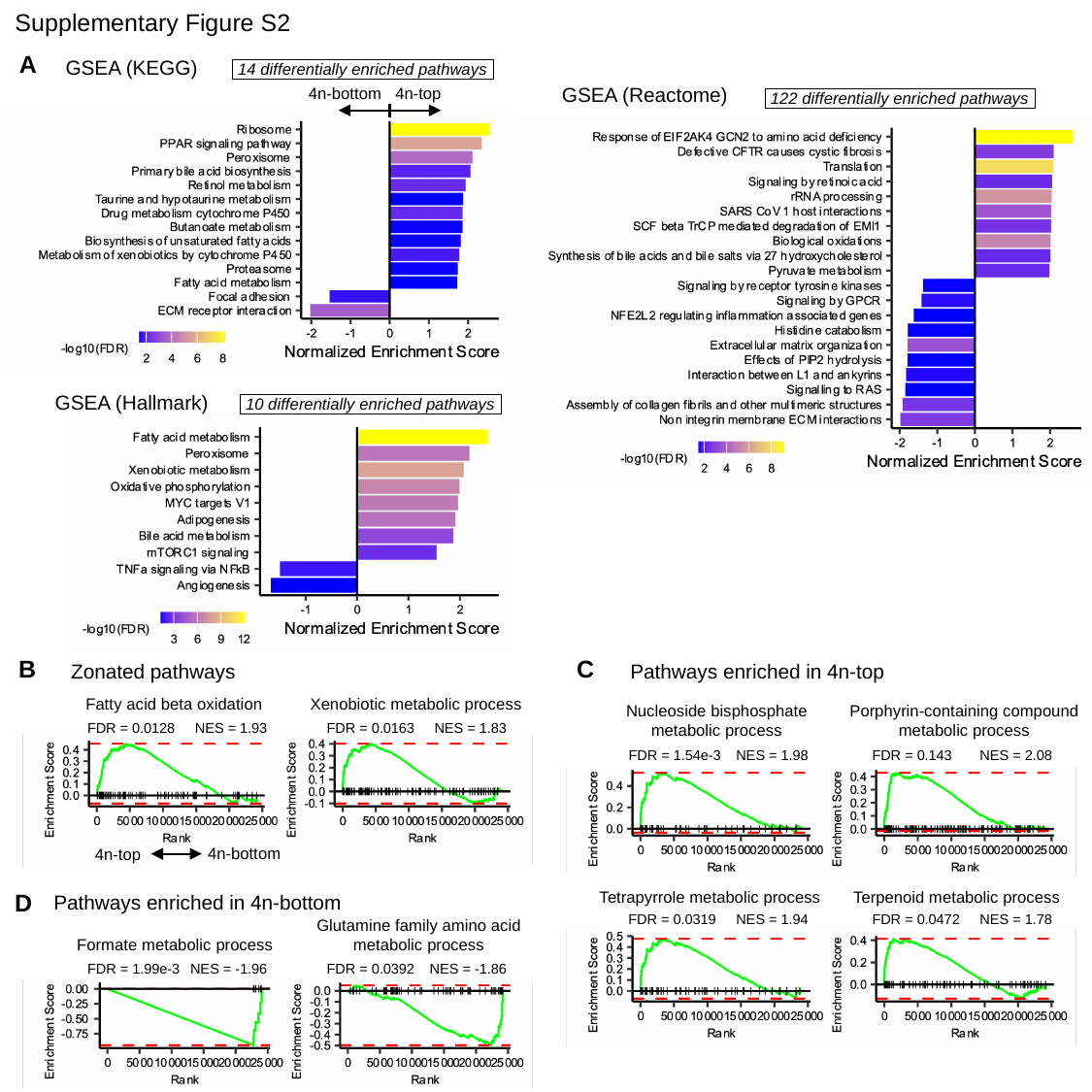

Supplementary Figure S2
A
GSEA (KEGG)
14 differentially enriched pathways
GSEA (Reactome)
4n-bottom 4n-top
122 differentially enriched pathways
GSEA (Hallmark)
10 differentially enriched pathways
B
C
 Zonated pathways
 Pathways enriched in 4n-top
Fatty acid beta oxidation
Xenobiotic metabolic process
Nucleoside bisphosphate metabolic process
Porphyrin-containing compound metabolic process
FDR = 0.0128
NES = 1.93
FDR = 0.0163
NES = 1.83
FDR = 1.54e-3
NES = 1.98
FDR = 0.143
NES = 2.08
4n-bottom
4n-top
Tetrapyrrole metabolic process
Terpenoid metabolic process
D
 Pathways enriched in 4n-bottom
FDR = 0.0319
NES = 1.94
FDR = 0.0472
NES = 1.78
Glutamine family amino acid metabolic process
Formate metabolic process
FDR = 1.99e-3
NES = -1.96
FDR = 0.0392
NES = -1.86
